# Contact-FP: a dimerization-dependent fluorescent protein toolkit for visualizing membrane contact site dynamics

**DOI:** 10.1101/2023.09.27.559820

**Authors:** Gregory E. Miner, Sidney Y. Smith, Wendy K. Showalter, Christina M. So, Joey V. Ragusa, Alex E. Powers, Maria Clara Zanellati, Chih-Hsuan Hsu, Michelle F. Marchan, Sarah Cohen

**Author notes:** Correspondences: Sarah Cohen.

## Abstract

Membrane contact sites (MCSs) are sites of close apposition between two organelles used to exchange ions, lipids, and information. Cells respond to changing environmental or developmental conditions by modulating the number, extent, or duration of MCSs. Because of their small size and dynamic nature, tools to study the dynamics of MCSs in live cells have been limited. Dimerization-dependent fluorescent proteins (ddFPs) targeted to organelle membranes are an ideal tool for studying MCS dynamics because they reversibly interact to fluoresce specifically at the interface between two organelles. Here, we build on previous work using ddFPs as sensors to visualize the morphology and dynamics of MCSs. We engineered a suite of ddFPs called Contact-FP that targets ddFP monomers to lipid droplets (LDs), the endoplasmic reticulum (ER), mitochondria, peroxisomes, lysosomes, plasma membrane, caveolae, and the cytoplasm. We show that these probes correctly localize to their target organelles. Using LDs as a test case, we demonstrate that Contact-FP pairs specifically localize to the interface between two target organelles. Titration of LD-mitochondria ddFPs revealed that these sensors can be used at high concentrations to drive MCSs or can be titrated down to minimally perturb and visualize endogenous MCSs. We show that Contact-FP probes can be used to: (1) visualize LD-mitochondria MCS dynamics, (2) observe changes in LD-mitochondria MCS dynamics upon overexpression of PLIN5, a known LD-mitochondrial tether, and (3) visualize two MCSs that share one organelle simultaneously (e.g., LD-mitochondria and LD-ER MCSs). Contact-FP probes can be optimized to visualize MCSs between any pair of organelles represented in the toolkit.

## Introduction

Eukaryotic cells are compartmentalized into membrane-bound organelles, allowing for the spatial separation of incompatible biochemical processes. Nevertheless, organelles must communicate for the cell to function as an integrated unit. One way that organelles communicate is at membrane contact sites (MCSs), sites of close apposition between the membranes of two organelles (Scorrano et al., 2019). Recent work has highlighted an increasing list of functions for MCSs, including roles in the exchange of ions, lipids, and proteins, as well as in organelle biogenesis and division (Prinz et al., 2019). Cells can modulate MCSs to adjust their metabolism in response to changing environmental or developmental conditions (Bohnert, 2020). Thus, there is a need for methods to visualize MCSs accurately and precisely in live cells. This has been challenging due to the small size and dynamic nature of these structures. Electron microscopy is considered the gold standard for identifying MCSs but is incompatible with live-cell imaging. Diffraction-limited fluorescence microscopy of cells expressing organelle markers can be used to measure organelle proximity and to infer changes in MCSs in response to different conditions (Valm et al., 2017). However, because MCSs are defined as occurring between membranes that are within a 30 nm distance (Scorrano et al., 2019), diffraction-limited microscopy cannot distinguish whether two organelles are in close proximity or truly form a *bone fide* MCS.

A variety of probes to detect MCSs have been developed to overcome these challenges. Bimolecular fluorescence complementation systems have been used to study multiple different MCSs (Jing et al., 2020). These systems involve targeting non-fluorescent fragments of fluorescent proteins (FPs) to two different organelles; when the organelles come together, the fluorophore is reconstituted, fluorescing only at the interface between the two membranes. Split GFP-based contact site (SPLICS) sensors have been developed for endoplasmic reticulum (ER)-mitochondria, ER-plasma membrane, ER-peroxisome, and mitochondria-peroxisome MCSs in mammalian cells (Cieri et al., 2017; Vallese et al., 2020). In *Saccharomyces cerevisiae*, split GFP-or split Venus-based probes have been used to investigate MCSs between nearly every pair of membrane-bound organelle (Kakimoto et al., 2018; Shai et al., 2018). These bimolecular fluorescence complementation systems are useful to detect differences in the number and extent of MCSs under different conditions, and in screens to identify new tethers at MCSs (Castro et al., 2022; Lahiri et al., 2014; Shai et al., 2018; Yang et al., 2018). However, because the fluorescence complementation of split fluorescent proteins is irreversible, these probes tend to stabilize MCSs and are therefore not ideal for studying MCS dynamics.

Dimerization-dependent fluorescent proteins (ddFPs) have emerged as an alternative tool for studying MCSs. ddFPs consist of two weakly fluorescent monomeric proteins that reversibly interact to form a fluorescent heterodimeric complex. The original ddFP was based on red fluorescent protein and was initially used to detect protein-protein interactions and caspase activity (Alford, Abdelfattah, et al., 2012). An extended palette in which green (GA) or red (RA) A monomers can be complemented interchangeably with a B monomer was subsequently developed (Alford, Ding, et al., 2012; Ding et al., 2015). Although ddFPs have primarily been used to study protein-protein interactions, the ddFP system has been adapted to detect MCSs including ER-mitochondria (Alford, Ding, et al., 2012; Naon et al., 2016; Nguyen & Voeltz, 2022), ER-P-body (Lee et al., 2020) or mitochondria-mitochondria (Abrisch et al., 2020) contacts. Because the interaction between ddFP monomers is reversible, ddFPs are more amenable than split FP systems for studying MCS dynamics. The main disadvantage of ddFPs is their lower fluorescence relative to split fluorescent proteins. However, we have found that with the availability of increasingly sensitive microscope detectors this disadvantage can now be overcome.

Here, we build on previous work using ddFPs to visualize the morphology and dynamics of MCSs. We engineered a suite of ddFPs called Contact-fluorescent protein (Contact-FP) that targets the three ddFP monomers (RA, GA, and B) to eight subcellular compartments: lipid droplets (LDs), ER, mitochondria, peroxisomes, lysosomes, plasma membrane, caveolae, and the cytoplasm. Using complementary cytoplasmic monomers (GA or B), we show that the seven organelle probes correctly localize to their targets. We demonstrate that Contact-FP pairs specifically localize to the interface between target organelles for each LD-organelle pair. We found that by titrating LD-mitochondria ddFPs, we could either induce MCSs at high concentrations, or minimally perturb and visualize endogenous MCSs at lower concentrations. Finally, we demonstrate several use cases for LD-mitochondria and LD-ER ddFPs.

Theoretically, Contact-FP probes can be optimized to visualize MCSs between any pair of organelles represented in the toolkit, although several optimization steps are required for each new ddFP pair.

## Results

### Design of Contact-FP Probes

Recent studies have demonstrated the use of ddFPs to visualize MCS dynamics (Abrisch et al., 2020; Alford, Ding, et al., 2012; Lee et al., 2020; Naon et al., 2016; Nguyen & Voeltz, 2022). Due to relatively low heterodimer affinity, the binding of ddFPs is reversible and therefore can be used to visualize MCS dynamics without artificially stabilizing them (Figure 1A). While ddFPs represent a promising tool for the study of MCS, constructs have only been made to target a small subset of organelle membranes. To expand the use of ddFPs for studying MCSs, we generated a ddFP construct library targeting ddFP monomers (GA, RA, and B) to the cytoplasm or to the cytoplasmic-facing membrane of seven organelles including the LD, ER, mitochondria, peroxisomes, lysosomes, plasma membrane, and caveolae. We named this suite of probes Contact-FP, analogous to Contact-ID, a split-pair BioID for identifying proteins at MCSs by proteomics (Kwak et al., 2020). Minimal membrane targeting domains were used to localize ddFPs when possible, to minimize disruption of organelle function (Figure 1B). To target ddFP monomers to the LD or peroxisome membrane we used the hairpin domain of GPAT4 (Wang et al., 2016) or the membrane targeting sequence of PEX3 (Soukupova et al., 1999)respectively, which we have successfully used previously to localize chimeric proteins (Miner et al., 2023). Monomers were targeted to the ER and mitochondrial membranes using the transmembrane domains of CYP2C1 and MAVS respectively (Cho et al., 2020). Plasma membrane targeted monomers were localized using the palmitoylation domain of GAP43 (Chung et al., 2019). Finally, due to a lack of well-established specific targeting domains, ddFP monomers were targeted to lysosomes and caveolae by fusing with lysosomal-associated membrane protein 1 (LAMP1) or caveolae-associated protein 1 (CAVIN1) respectively.

**Figure 1.**
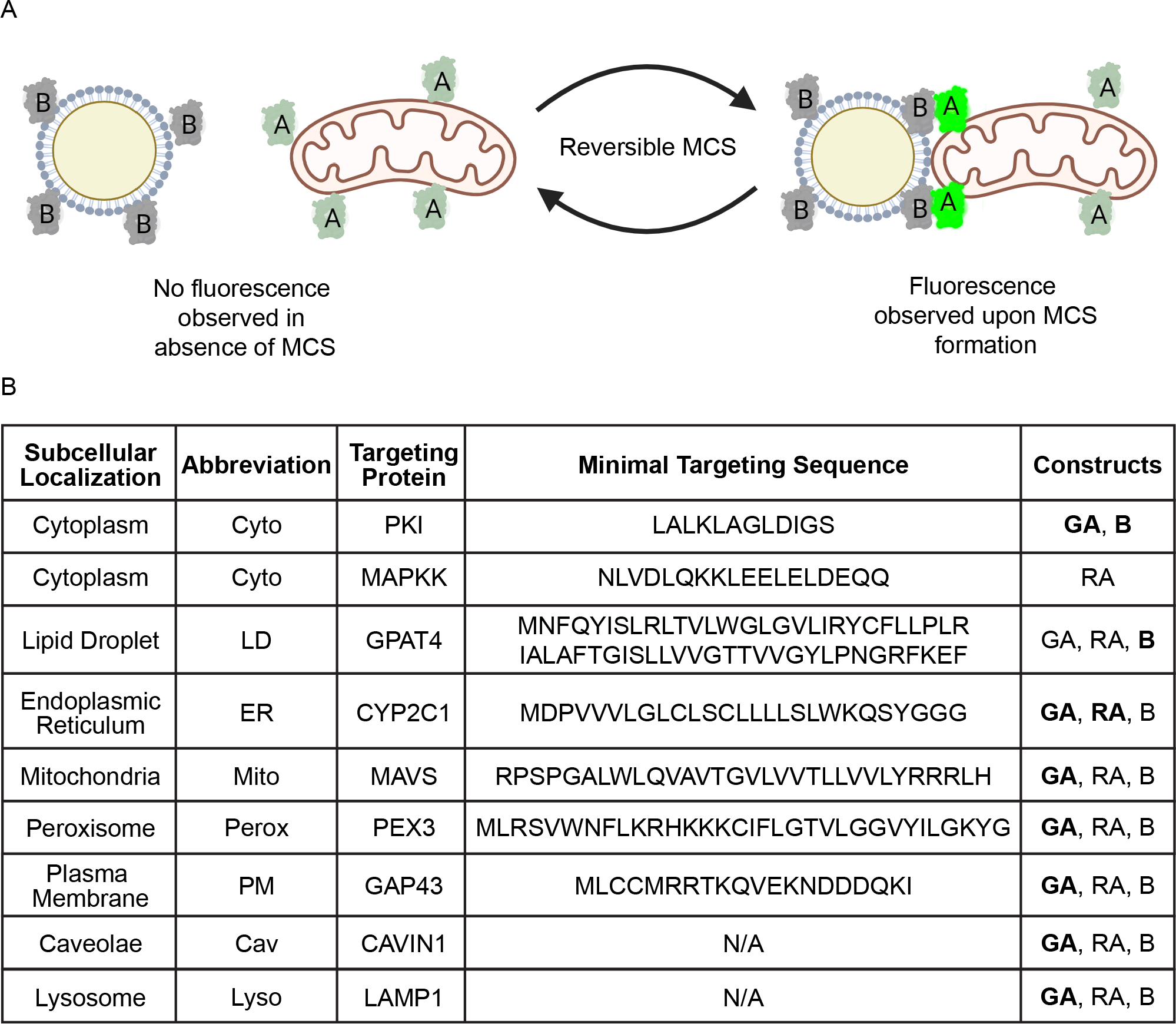
Design of Contact-FP Probes. **(A)** Cartoon of reversible Contact-FP probe system. Monomers targeted to distinct cellular membranes reversibly dimerize when in close proximity (∼10-30 nM) resulting in increased fluorescence of the “A” monomer at membrane contact sites. Cartoon created with BioRender.com **(B)** List of probes generated for the Contact-FP library. The targeting sequence used to localize probes and the source protein are included. Probes demonstrated within this study are bolded.

### Contact-FP Probes Localize to their Target Organelles

We next tested whether our organelle-targeted monomers properly localized to the cytoplasmic face of the intended organelle membrane by co-expressing them with the cytoplasmic heterodimeric partner (Figure 2A). For these experiments we used the GA monomer constructs as these exhibit higher brightness and contrast than the RA monomer probes, and therefore were ideal for initial optimization and evaluation of novel ddFP constructs. We found that co-expression of cytoplasmic monomer with our organelle-anchored monomers resulted in fluorescence signal along the entire surface of the organelle of interest (Figure 2B-H). Localization was confirmed by comparing localization with known fluorescent organelle markers by confocal microscopy. Given that ER proteins can be restricted to either ER tubules or sheets, we utilized Airyscan superresolution imaging to resolve both substructures and confirmed localization of ER-tethered GA (ER-GA) along the entire ER surface. While most constructs labeled the target organelle following a 24-hr transfection, LD-targeted constructs required 48 hrs for efficient labeling of LDs. Unlabeled LDs likely represent mature LDs detached from the ER which cannot be labeled by membrane anchored proteins that require ER-LD bridges for trafficking (Wilfling et al., 2013).

**Figure 2.**
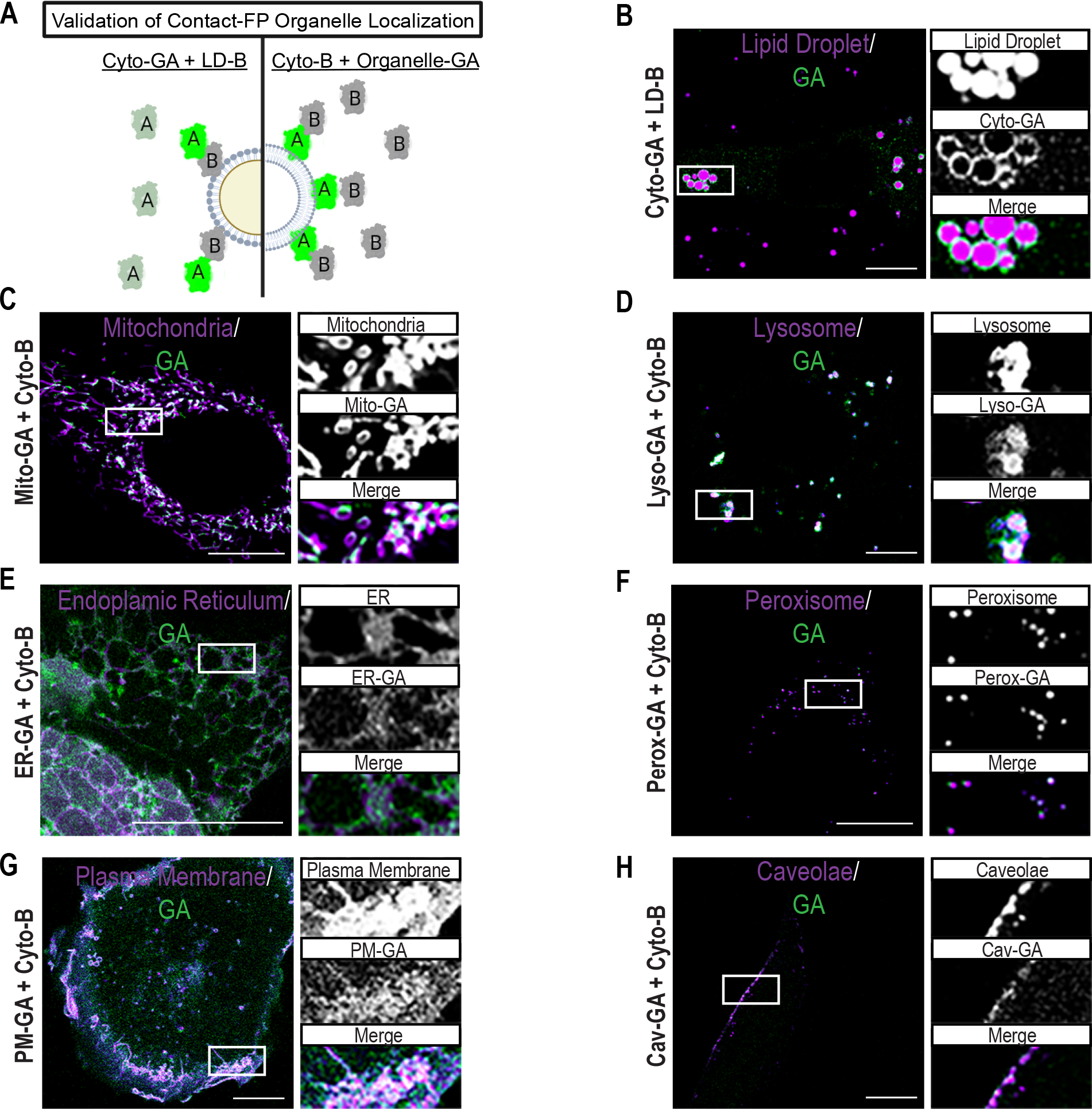
Contact-FP Probes Localize to their Target Organelles. **(A)** Cartoon depicting use of Cytoplasmic Contact-FP probes (Cyto-GA and Cyto-B) to validate cellular localization of organelle membrane targeted probes. Cartoon created with BioRender.com **(B)** Single focal plane micrographs of U-2 OS cells transfected with the indicated Contact-FP probes and labeled for LDs with Bodipy 665/676. Merged micrograph includes GA and Bodipy 665/667 channels to demonstrate co-localization (white). **(C-H)** Single focal plane micrographs of U-2 OS cells transfected with the indicated Contact-FP probes and labeled for organelles with (C) mApple-TOMM20-N-10, (D) mApple-Lysosomes-20, (E) mApple-Sec61B, (F) mApple-Peroxisomes-2, (G) mApple-Farnesyl-5, (H) mApple-Caveolin-C-10). Merged micrograph includes GA and mApple channels to demonstrate co-localization (white). (**C**,**E**,**F**) U-2 OS cells were imaged using Airyscan. Scale bar, 10μm.

### LD-Organelle Contact-FP Pairs Correctly Localize to MCSs

Having confirmed that the Contact-FP constructs properly localize, we next validated their use to specifically detect MCSs. For these experiments we focused on studying LD-organelle MCSs. LDs are an ideal use of Contact-FP as they form dynamic and transient MCSs with all organelles included in our set of constructs (Matthaeus & Taraska, 2021; Valm et al., 2017). Specifically, we tested whether organelle-tethered GA monomers would brightly fluoresce when co-localized with a LD-tethered B monomer (LD-B) at LD-organelle MCSs (Figure 3A). As expected, we were able to observe LD-organelle contacts with all organelles tested and found that GA fluorescence was brightest at sites of organelle marker co-localization (Figure 3 B-G). It was necessary to optimize the ratio of GA and B probes for each pair of organelles. Factors that could affect the amount of probe needed include the specific turnover rate of each probe, and the abundance of the MCS being studied.

**Figure 3.**
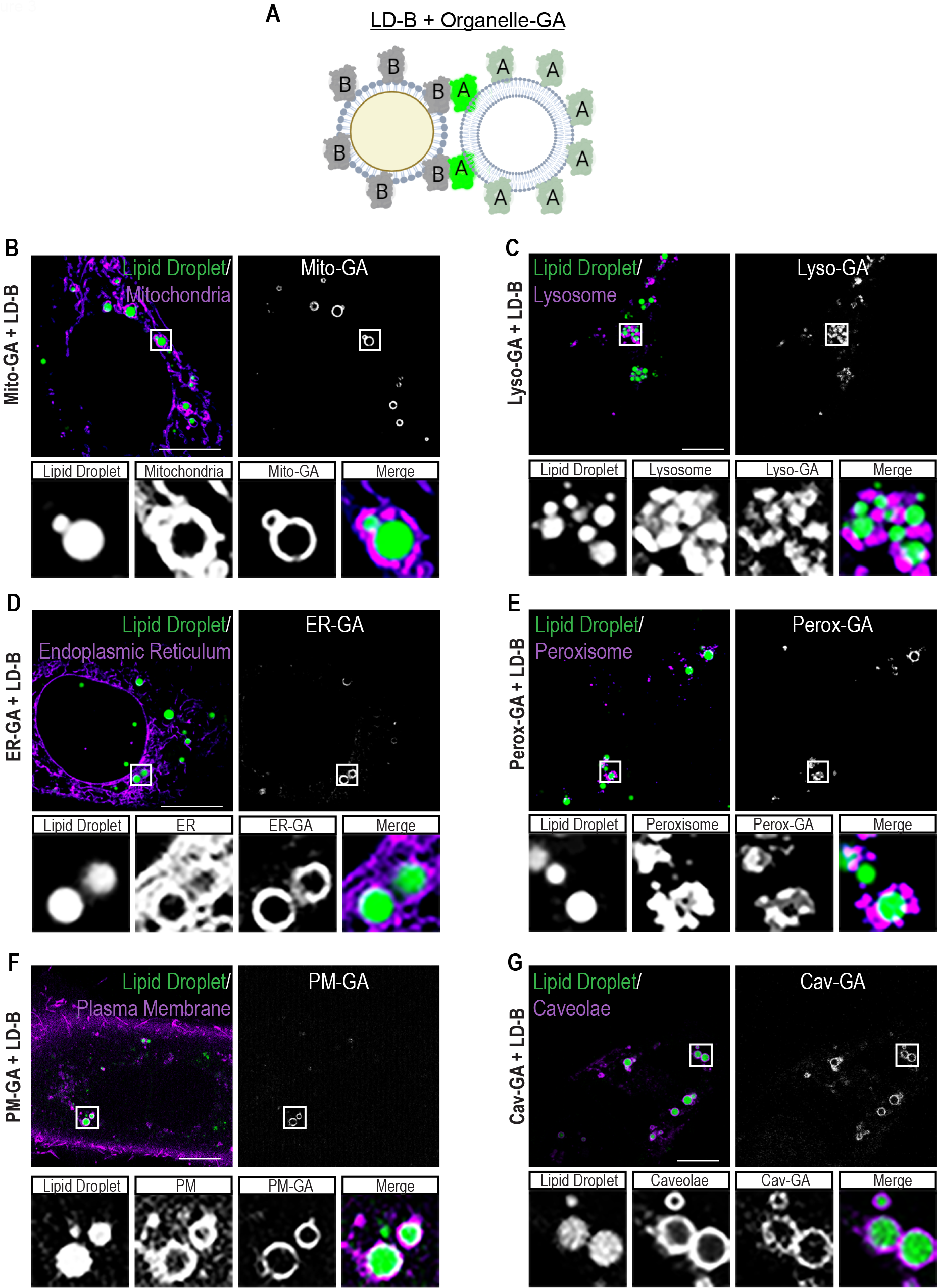
LD-Organelle Contact-FP Pairs Correctly Localize to MCSs. **(A)** Cartoon depicting use of Contact-FP probes (organelle-GA and LD-B) to validate Contact-FP fluorescence at MCSs. Cartoon created with BioRender.com **(B-G)** Single focal plane micrographs of U-2 OS cells transfected with the indicated Contact-FP probes, labeled for LDs with Bodipy 665/676, and labeled for organelles with (B) mApple-TOMM20-N-10, (C) mApple-Lysosomes-20, (D) mApple-Sec61B, (E) mApple-Peroxisomes-2, (F) mApple-Farnesyl-5, (G) mApple-Caveolin-C-10. Merged micrograph includes Bodipy 665/676 and mApple channels (but not GA) to demonstrate LD-organelle MCSs (white). (**B**,**D**,**E**) U-2 OS cells were imaged using Airyscan. Scale bar, 10μm.

The abundance of MCSs varied greatly depending on the LD-organelle pair. For example, most LDs in cells transfected with mitochondria-targeted GA (mito-GA) and LD-B were positive for GA signal (Figure 3B), indicating that LD-mitochondria contacts are abundant in U-2 OS cells. In contrast, GA signal was only observed on LDs at the cell periphery in cells expressing plasma membrane-targeted GA (PM-GA) and LD-B (Figure 3F), indicating that only a subset of LDs form MCSs with the plasma membrane in U-2 OS cells. Qualitatively, organelle morphology did not appear disrupted by the expression of ddFP probes. The exception was that upon expression of peroxisome-targeted GA (perox-GA) and LD-B, rings of peroxisomes were observed around LDs in cells with detectable GA signal (Figure 3E), while GA signal was not detectable in the majority of transfected cells. Rings of peroxisomes were never observed in control cells. We interpret this to mean that LD-peroxisome contacts are exceedingly rare in U-2 OS cells. Thus, the only cells in which we detected GA signal were those in which ddFPs were expressed at concentrations high enough to induce the formation of MCSs.

### LD-mitochondria Contact-FP Probes can be Titrated to Induce or Visualize MCSs

Given the low affinity of the heterodimeric monomers and therefore reversible binding, we hypothesized that induced MCS formation (e.g., in the case of LD-peroxisome MCSs) was a consequence of high Contact-FP expression. A major goal of our Contact-FP probes is the ability to monitor MCS dynamics while not perturbing MCS formation. Therefore, we tested whether the expression of Contact-FP pairs could be titrated to avoid MCS induction while retaining robust signal. We expressed varying amounts of mito-GA together with a constant level of LD-B (Figure 4). We found that expressing high concentrations of mito-GA caused a significant increase in LD-mito colocalization and a corresponding increase in GA fluorescence at LDs relative to controls expressing only a single monomer (Figure 4A-B, and 4E-F). In contrast, reduced expression of mito-GA did not significantly induce LD-mito contacts, while strong GA signal was retained at contact sites (Figure 4C, and 4E-F). Further reduction of mito-GA expression resulted in visible yet faint GA signal at the LD-mito MCS (Figure 4D-F). Taken together, we demonstrate that expression of Contact-FP probes can be used to either induce MCSs at high concentrations or to visualize MCS dynamics without artificial contact site induction at low concentrations.

**Figure 4.**
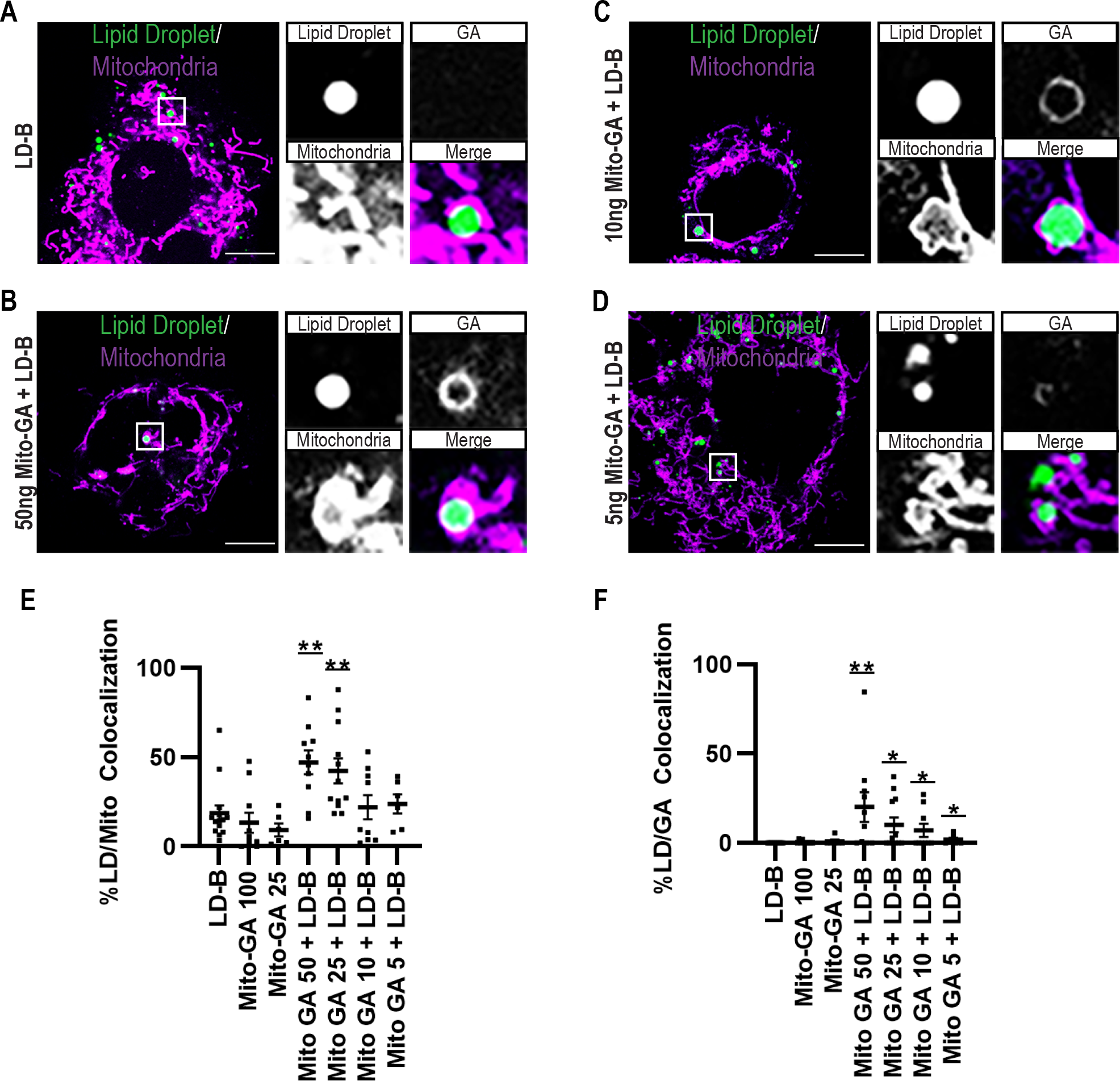
LD-mitochondria Contact-FP Probes can be Titrated to Induce or Visualize MCSs. **(A-D)** Single focal plane micrographs of U-2 OS cells transfected with the indicated Contact-FP probes, labeled for LDs with Bodipy 665/676, and labeled for mitochondria with mApple-TOMM20-N-10. Merged micrograph includes Bodipy 665/676 and mApple channels (but not GA) to demonstrate LD-organelle MCSs (white). **(E-F)** Quantification of images from (A-D). (E) Colocalization of LDs with mitochondria was measured as % LD pixels overlapping with mitochondria, and (F) Colocalization of LDs with Mito-GA was measured as % LD pixels overlapping with Mito-GA. N=3 biological replicates. Scale bar, 10μm. Error bars represent ± SEM. *p < 0.05, **p < 0.01.

### Use Cases for Contact-FP Probes: LD-mitochondria Dynamics in Response to Overexpression of Known Tethers and Visualizing Multiple MCSs

Our data thus far demonstrate that Contact-FP probes properly localize to their target organelle membranes and can be used to visualize MCSs without MCS induction. Finally, we wanted to demonstrate potential uses of our Contact-FP probes. We first tested the use of our LD-mito Contact-FP probes to investigate the effect of PLIN5 overexpression on LD-mito MCS dynamics (Figure 5). PLIN5 is a well-established LD-mito tether that has been shown to induce membrane contacts upon overexpression and reduce LD mobility (Miner et al., 2023; Wang et al., 2011). As anticipated, we observed a drastic increase in LD-mito contacts following overexpression of PLIN5 and an increase in GA fluorescence relative to controls (Figure 5A-B). Intriguingly, time-lapse imaging showed that LD-mito MCSs in control cells are highly dynamic and typically only cover a portion of the LD surface. Over time we observed rapid changes in MCS area at individual LDs, with LD-mito contact sites growing and shrinking (Figure 5C). In contrast, LDs in cells expressing PLIN5 appear enveloped by mitochondria with extensive LD-mito MCSs. Additionally, these MCSs appear extremely stable and no longer change in size over time (Figure 5D). This example illustrates the ability to use Contact-FP to directly visualize MCS dynamics.

**Figure 5.**
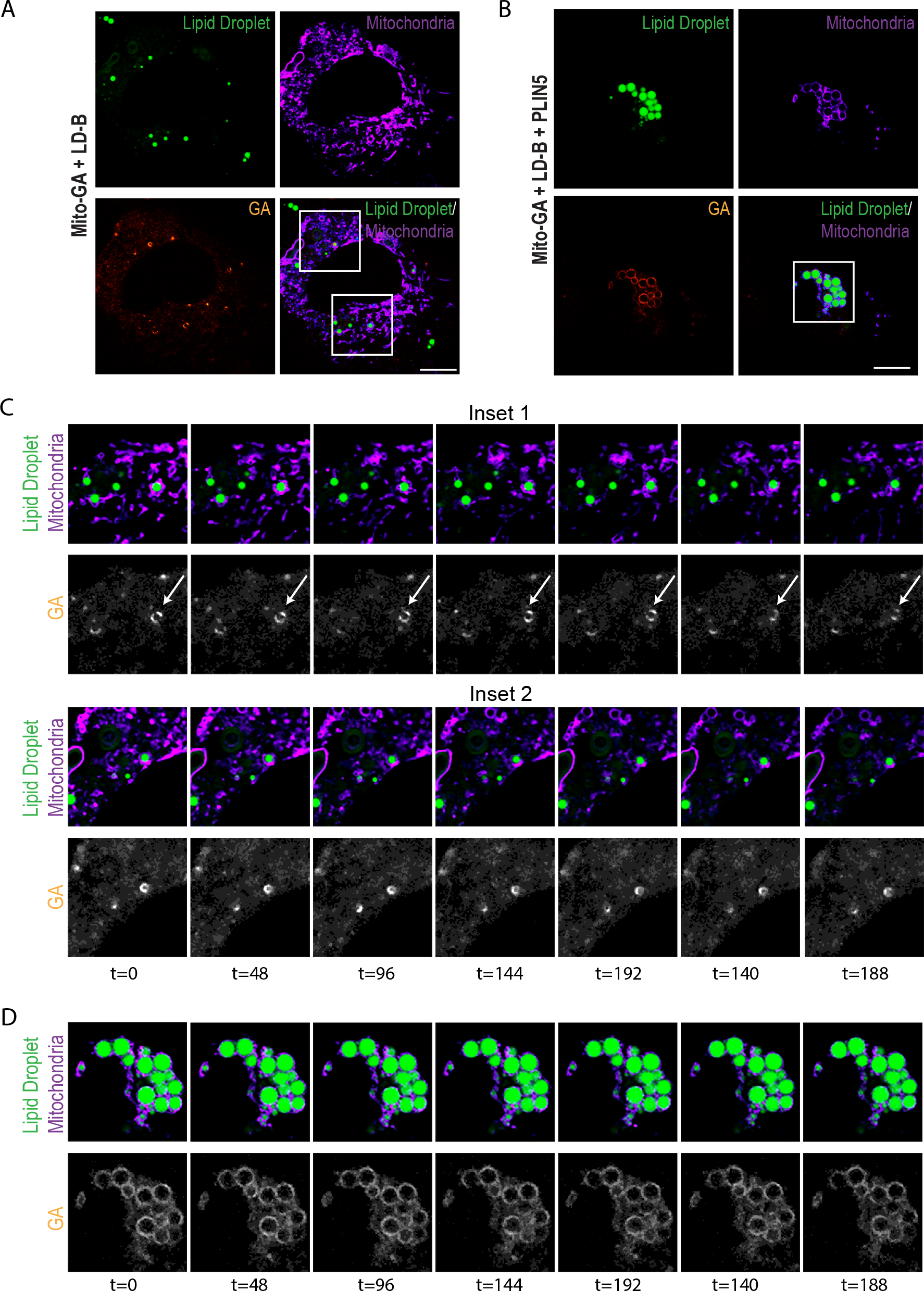
Use Case for Contact-FP Probes: LD-mitochondria Dynamics in Response to Overexpression of Known Tether. **(A-B)** Single focal plane micrographs of U-2 OS cells transfected with the indicated constructs, labeled for LDs with Bodipy 665/676, and labeled for mitochondria with mApple-TOMM20-N-10. Merged micrograph includes Bodipy 665/676 and mApple channels to demonstrate LD-organelle MCSs (white). **(C)** Insets show dynamics of LD-mitochondria MCSs over a 3 min time-lapse movie with frames captured every 12 sec. White arrow indicates a LD-mitochondria MCS which dissipates during the movie. **(D)** Inset shows decreased dynamics of LD-mitochondria MCSs in cells overexpressing the LD-mitochondria tether PLIN5 over a 3 min time-lapse movie with frames captured every 12 sec. Scale bar, 10μm.

Another advantage of Contact-FP is the ability to choose from constructs containing a fluorescent GA or RA monomer, which fluoresce at green and red wavelengths respectively. Since both GA and RA monomers can form heterodimeric pairs with the B monomer, this opens the possibility to monitor two MCSs simultaneously. As the LD is known to interact extensively with both mitochondria and the ER, we assessed whether we could visualize both LD-mito and LD-ER MCSs in the same cell (Figure 6A). We found that co-expression of Mito-GA and ER-RA with LD-B allowed simultaneous visualization of both LD-mito and LD-ER MCSs (Figure 6B-C). As expected, our GA and RA signal colocalizes with LDs (Figure 6B) and respective organelle membranes (Figure 6C). Further, RA and GA signals are spatially distinct with no significant crosstalk. Intriguingly, while most LDs appear to form MCSs with either mitochondria or ER, we also observed cases where a single LD forms three-way MCSs with both mitochondria and ER (Figure 6C, inset).

**Figure 6.**
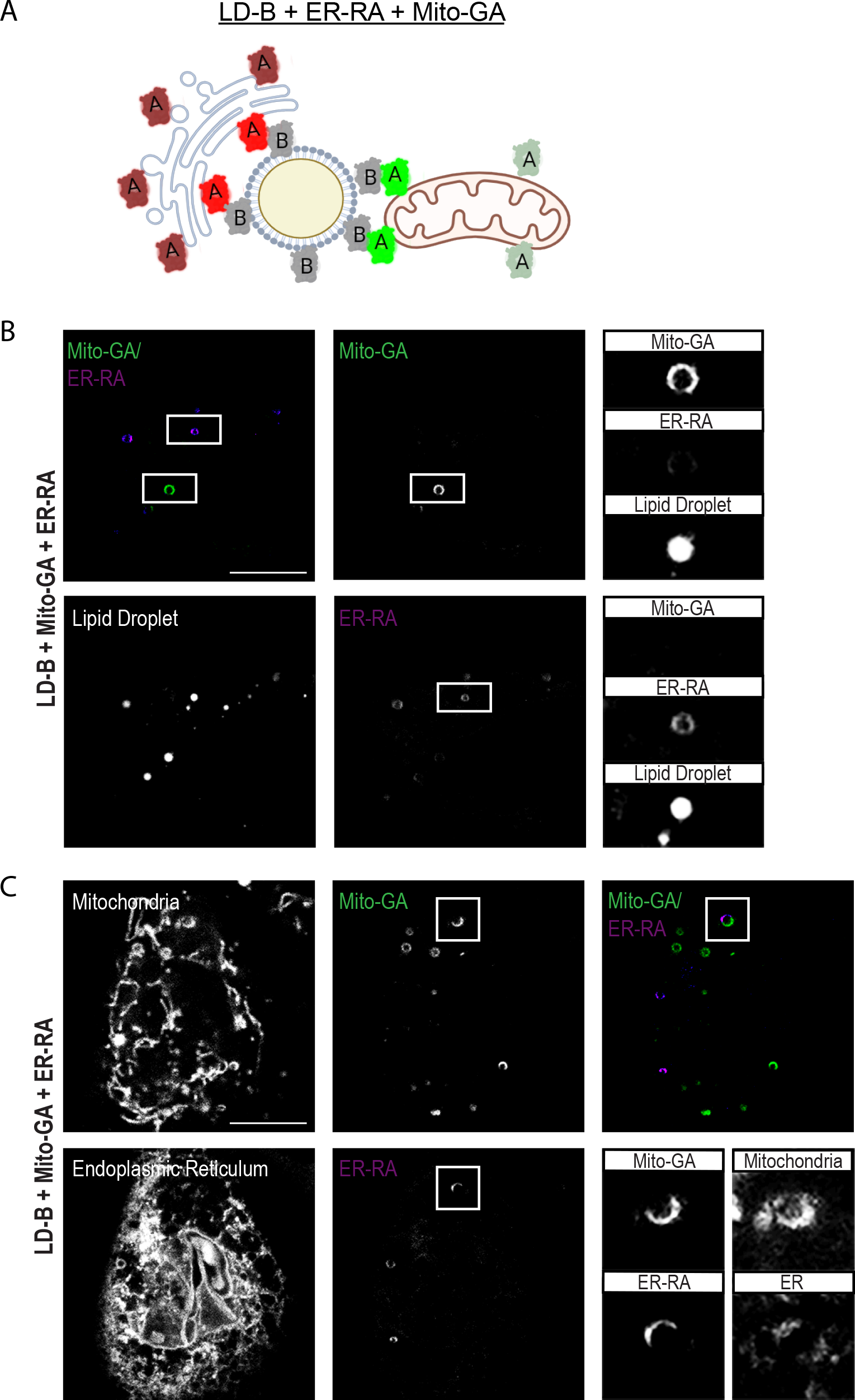
Use Case for Contact-FP Probes: Visualizing Multiple MCSs. **(A)** Cartoon depicting use of Contact-FP probes (Mito-GA, ER-RA, and LD-B) to simultaneously visualize LD-mitochondria and LD-ER MCSs. Cartoon created with BioRender.com **(B)** Single focal plane micrographs of U-2 OS cells transfected with the indicated constructs and labeled for LDs with Lipi-blue. Merged micrograph includes GA and RA channels to demonstrate Mito and ER MCSs are spatially distinct. **(C)** Single focal plane micrographs of U-2 OS cells transfected with the indicated constructs, labeled for mitochondria with MitoTracker Deep Red and labeled for ER with TagBFP-KDEL. Merged micrograph includes GA and RA channels to demonstrate Mito and ER MCSs are spatially distinct. Scale bar, 10μm.

## Discussion

Here, we have built on previous work using ddFPs as sensors to visualize MCSs. We engineered a suite of ddFPs called Contact-FP that targets ddFP monomers to seven organelles or the cytoplasm. We show that these probes correctly localize to their target organelles and that Contact-FP pairs specifically localize to the interface between target organelles. Contact-FP can be used at high concentrations to drive MCSs or can be titrated down to minimally perturb and visualize endogenous MCSs. We demonstrated several use cases for Contact-FP probes, including the observation of changes in LD-mitochondria MCS dynamics upon overexpression of PLIN5, and the visualization of two MCSs that share one organelle simultaneously.

Contact-FP probes can be used to visualize MCSs between any pair of organelles represented in the toolkit. However, optimization is required for each organelle pair, and is cell-type dependent. We recommend performing titration experiments such as in Figure 4 to determine the minimal expression of a Contact-FP pair that provides robust signal without inducing MCSs or disrupting organelle morphology in a given cell type. Making stable cell lines or introducing ddFPs via CRISPR may provide the most consistent cell-to-cell expression levels of ddFPs. The main advantage of ddFPs over bimolecular fluorescence complementation systems such as split-GFP is that ddFP dimerization is reversible, allowing the visualization of MCS dynamics with minimal perturbation of the endogenous MCS. However, the disadvantage is that ddFPs are not as bright as split-FPs. For most of the LD-organelle MCSs we imaged, we found that confocal microscopy was sufficiently sensitive to detect GA signal at contact sites. However, several MCSs benefitted from more sensitive methods such as Airyscan microscopy. We envision that Contact-FP will prove useful for characterizing MCSs in many contexts.

It will be fascinating to catalogue the abundance and dynamics of MCSs in different cell types, which can have highly specialized organelle morphologies and dynamics. MCSs can remodel in response to developmental or environmental cues (Bohnert, 2020; Prinz et al., 2019). We anticipate that Contact-FP will provide insight into how MCS dynamics change throughout processes such as cell division, migration, and differentiation. Because Contact-FP works on short timescales, it will also be useful for measuring changes in MCS dynamics in response to acute changes such as infection, nutritional fluctuations, and pharmacological perturbations. Finally, it may eventually be possible to introduce Contact-FP probes *in vivo*, allowing minimally perturbing characterization of MCSs in the context of developing and aging organisms, and in various models of human diseases.

### Resource Availability

#### Lead Contact

Further information and requests about resources and reagents are available from the corresponding author, Sarah Cohen, upon reasonable request.

#### Materials availability

Plasmids generated in this study will be made available from the lead contact on request.

#### Data availability

Any additional information required to reanalyze the data reported in this paper is available from the lead contact upon reasonable request.

### Experimental model and subject details

U-2 OS cells were obtained from the UNC Tissue Culture Facility and maintained in Dulbecco’s Modified Eagle Medium (DMEM) with 10% FBS and 4 mM glutamine (complete medium, CM). Cells were cultured on chambered cover glass (#1.5 high performance cover glass, Cellvis), coated with 10 μg/ml fibronectin (MilliporeSigma).

### Method details

#### Cell transfection

Cells were transfected with mApple-TOMM20-N-10, mApple-Lysosomes-20, mApple-Sec61B, mApple-Peroxisomes-2, mApple-Farnesyl-5, Apple-Caveolin-C-10, TagBFP-KDEL or Contact-FP constructs using Lipofectamine 2000 (Invitrogen), according to the manufacturer’s instructions. All experiments were performed in 8-well chambered cover glass (Cellvis). Cells were transected with the following range of plasmid amounts: Cyto-B/Cyto-GA (200-50 ng), LD-B (50-10 ng), Organelle-GA (100-10 ng). Cells were imaged either 24 hrs following transfection, or 48 hrs following transfection if LD-B was used.

#### Plasmids

Contact-FP constructs were generated using HiFi DNA Assembly Master mix (New England Biolabs, E2621). For plasmid construction, all PCRs were performed using Q5 High Fidelity DNA polymerase (M0419; New England Biolabs) and restriction enzymes from New England Biolabs. The following plasmids were kind gifts: RA-NES, GA-NES, and GB-NES from Robert Campbell (University of Alberta) (Ding et al., 2015), TagBFP-KDEL, RA-Sec61B, GA-MFF, and GB-Dcp1b from Gia Voeltz (University of Colorado) (Friedman et al., 2011; Lee et al., 2020), Apex2-OMM and ERM-Apex2 from Alice Ting (Stanford) (Lam et al., 2014), PM-FRB-CFP from Tamas Balla (NICHHD) (Varnai et al., 2006), Lamp1-mTurquoise2 from Dorus Gadella (University of Amsterdam) (Chertkova et al., 2020, BioRxiv, 160374), cavin-1-mEGFP from Ari Helenius (ETH Zurich) (Hayer et al., 2010), mApple-TOMM20-N-10, mApple-Lysosomes-20, mApple-Peroxisomes-2, mApple-Farnesyl-5, and mApple-Caveolin-C-10 from Michael Davidson (Florida State University), and mApple-Sec61B from Jennifer Lippincott-Schwartz (Janelia Research Campus) (Nixon-Abell et al., 2016).

#### Chemicals and Dyes

The following chemicals and dyes were used: 10 μg/ml fibronectin (MilliporeSigma), 50 ng/ml BODIPY 665/676 (Life Technologies), 100 nM MitoTracker Deep Red (Life Technologies), 50 nM Lipi-Blue (Dojindo).

#### Microscopy and image processing

Images were acquired on an inverted Zeiss 800/Airyscan laser scanning confocal microscope equipped with 405, 488, 561 and 647 nm diode lasers, and Galium Arsenide Phosphid (GaAsP) and Airyscan detectors. Confocal and Airyscan images were acquired using a 63x/1.4 NA objective lens, at 37 °C and 5% CO_2_ (Carl Zeiss, Oberkochen, Germany). Airyscan images were processed in Zen software (Carl Zeiss) using a processing strength of 6.0. Image brightness and contrast were adjusted in Adobe Photoshop CS.

### Quantification and statistical analysis

Images were analyzed using CellProfiler (Stirling et al., 2021). For colocalization, masks of LD, mitochondria, and GA were created using the corresponding channels. For overlap of LDs with mitochondria and GA, the percentage of LD pixels colocalized with the Mitochondria or GA versus total LD pixels was calculated. Statistical detail for all experiments can be found in the figure legends. Statistical analysis among groups was performed using Student’s t test.

## References

Abrisch, R. G., Gumbin, S. C., Wisniewski, B. T., Lackner, L. L., & Voeltz, G. K. (2020). Fission and fusion machineries converge at ER contact sites to regulate mitochondrial morphology. Journal of Cell Biology, 219(4). 10.1083/JCB.201911122/VIDEO-9

Alford, S. C., Abdelfattah, A. S., Ding, Y., & Campbell, R. E. (2012). A Fluorogenic Red Fluorescent Protein Heterodimer. Chemistry & Biology, 19(3), 353–360. 10.1016/J.CHEMBIOL.2012.01.006

Alford, S. C., Ding, Y., Simmen, T., & Campbell, R. E. (2012). Dimerization-dependent green and yellow fluorescent proteins. ACS Synthetic Biology, 1(12), 569–575. 10.1021/SB300050J

Bohnert, M. (2020). Tether Me, Tether Me Not—Dynamic Organelle Contact Sites in Metabolic Rewiring. Developmental Cell, 54(2), 212–225. 10.1016/J.DEVCEL.2020.06.026

Castro, I. G., Shortill, S. P., Dziurdzik, S. K., Cadou, A., Ganesan, S., Valenti, R., David, Y., Davey, M., Mattes, C., Thomas, F. B., Avraham, R. E., Meyer, H., Fadel, A., Fenech, E. J., Ernst, R., Zaremberg, V., Levine, T. P., Stefan, C., Conibear, E., & Schuldiner, M. (2022). Systematic analysis of membrane contact sites in Saccharomyces cerevisiae uncovers modulators of cellular lipid distribution. ELife, 11. 10.7554/ELIFE.74602

Cho, K. F., Branon, T. C., Rajeev, S., Svinkina, T., Udeshi, N. D., Thoudam, T., Kwak, C., Rhee, H. W., Lee, I. K., Carr, S. A., & Ting, A. Y. (2020). Split-TurboID enables contact-dependent proximity labeling in cells. Proceedings of the National Academy of Sciences of the United States of America, 117(22), 12143. 10.1073/PNAS.1919528117/-/DCSUPPLEMENTAL

Chung, J., Wu, X., Lambert, T. J., Lai, Z. W., Walther, T. C., & Farese, R. V. (2019). Lipid droplet assembly factor-1 and seipin form a lipid droplet assembly complex. Developmental Cell, 51(5), 551. 10.1016/J.DEVCEL.2019.10.006

Cieri, D., Vicario, M., Giacomello, M., Vallese, F., Filadi, R., Wagner, T., Pozzan, T., Pizzo, P., Scorrano, L., Brini, M., & Calí, T. (2017). SPLICS: a split green fluorescent protein-based contact site sensor for narrow and wide heterotypic organelle juxtaposition. Cell Death & Differentiation 2018 25:6, 25(6), 1131–1145. 10.1038/s41418-017-0033-z

Ding, Y., Li, J., Enterina, J. R., Shen, Y., Zhang, I., Tewson, P. H., Mo, G. C. H., Zhang, J., Quinn, A. M., Hughes, T. E., Maysinger, D., Alford, S. C., Zhang, Y., & Campbell, R. E. (2015). Ratiometric biosensors based on dimerization-dependent fluorescent protein exchange. Nature Methods 2015 12:3, 12(3), 195–198. 10.1038/nmeth.3261

Friedman, J. R., Lackner, L. L., West, M., DiBenedetto, J. R., Nunnari, J., & Voeltz, G. K. (2011). ER tubules mark sites of mitochondrial division. Science, 334(6054), 358–362. 10.1126/SCIENCE.1207385/SUPPL_FILE/FRIEDMAN.SOM.REV1.PDF

Hayer, A., Stoeber, M., Bissig, C., & Helenius, A. (2010). Biogenesis of Caveolae: Stepwise Assembly of Large Caveolin and Cavin Complexes. Traffic, 11(3), 361–382. 10.1111/J.1600-0854.2009.01023.X

Jing, J., Liu, G., Huang, Y., & Zhou, Y. (2020). A molecular toolbox for interrogation of membrane contact sites. The Journal of Physiology, 598(9), 1725. 10.1113/JP277761

Kakimoto, Y., Tashiro, S., Kojima, R., Morozumi, Y., Endo, T., & Tamura, Y. (2018). Visualizing multiple inter-organelle contact sites using the organelle-targeted split-GFP system. Scientific Reports 2018 8:1, 8(1), 1–13. 10.1038/s41598-018-24466-0

Kwak, C., Shin, S., Park, J. S., Jung, M., My Nhung, T. T., Kang, M. G., Lee, C., Kwon, T. H., Park, S. K., Mun, J. Y., Kim, J. S., & Rhee, H. W. (2020). Contact-ID, a tool for profiling organelle contact sites, reveals regulatory proteins of mitochondrial-associated membrane formation. Proceedings of the National Academy of Sciences of the United States of America, 117(22). 10.1073/pnas.1916584117

Lahiri, S., Chao, J. T., Tavassoli, S., Wong, A. K. O., Choudhary, V., Young, B. P., Loewen, C. J. R., & Prinz, W. A. (2014). A Conserved Endoplasmic Reticulum Membrane Protein Complex (EMC) Facilitates Phospholipid Transfer from the ER to Mitochondria. PLOS Biology, 12(10), e1001969. 10.1371/JOURNAL.PBIO.1001969

Lam, S. S., Martell, J. D., Kamer, K. J., Deerinck, T. J., Ellisman, M. H., Mootha, V. K., & Ting, A. Y. (2014). Directed evolution of APEX2 for electron microscopy and proximity labeling. Nature Methods 2014 12:1, 12(1), 51–54. 10.1038/nmeth.3179

Lee, J. E., Cathey, P. I., Wu, H., Parker, R., & Voeltz, G. K. (2020). Endoplasmic reticulum contact sites regulate the dynamics of membraneless organelles. Science, 67(6477). 10.1126/SCIENCE.AAY7108/SUPPL_FILE/AAY7108_LEE_SM.PDF

Matthaeus, C., & Taraska, J. W. (2021). Energy and Dynamics of Caveolae Trafficking. Frontiers in Cell and Developmental Biology, 8, 614472. 10.3389/FCELL.2020.614472/BIBTEX

Miner, G. E., So, C. M., Edwards, W., Ragusa, J. V., Wine, J. T., Wong Gutierrez, D., Airola, M. V., Herring, L. E., Coleman, R. A., Klett, E. L., & Cohen, S. (2023). PLIN5 interacts with FATP4 at membrane contact sites to promote lipid droplet-to-mitochondria fatty acid transport. Developmental Cell, 58(14), 1250–1265.e6. 10.1016/J.DEVCEL.2023.05.006

Naon, D., Zaninello, M., Giacomello, M., Varanita, T., Grespi, F., Lakshminaranayan, S., Serafini, A., Semenzato, M., Herkenne, S., Hernández-Alvarez, M. I., Zorzano, A., De Stefani, D., Dorn, G. W., & Scorrano, L. (2016). Critical reappraisal confirms that Mitofusin 2 is an endoplasmic reticulum-mitochondria tether. Proceedings of the National Academy of Sciences of the United States of America, 113(40), 11249–11254. 10.1073/PNAS.1606786113/SUPPL_FILE/PNAS.201606786SI.PDF

Nguyen, T. T., & Voeltz, G. K. (2022). An ER phospholipid hydrolase drives ER-associated mitochondrial constriction for fission and fusion. ELife, 11. 10.7554/ELIFE.84279

Nixon-Abell, J., Obara, C. J., Weigel, A. V., Li, D., Legant, W. R., Xu, C. S., Pasolli, H. A., Harvey, K., Hess, H. F., Betzig, E., Blackstone, C., & Lippincott-Schwartz, J. (2016). Increased spatiotemporal resolution reveals highly dynamic dense tubular matrices in the peripheral ER. Science, 354(6311). 10.1126/SCIENCE.AAF3928/SUPPL_FILE/NIXON-ABELL.SM.PDF

Prinz, W. A., Toulmay, A., & Balla, T. (2019). The functional universe of membrane contact sites. Nature Reviews Molecular Cell Biology 2019 21:1, 21(1), 7–24. 10.1038/s41580-019-0180-9

Scorrano, L., De Matteis, M. A., Emr, S., Giordano, F., Hajnóczky, G., Kornmann, B., Lackner, L. L., Levine, T. P., Pellegrini, L., Reinisch, K., Rizzuto, R., Simmen, T., Stenmark, H., Ungermann, C., & Schuldiner, M. (2019). Coming together to define membrane contact sites. Nature Communications 2019 10:1, 10(1), 1–11. 10.1038/s41467-019-09253-3

Shai, N., Yifrach, E., Van Roermund, C. W. T., Cohen, N., Bibi, C., Ijlst, L., Cavellini, L., Meurisse, J., Schuster, R., Zada, L., Mari, M. C., Reggiori, F. M., Hughes, A. L., Escobar-Henriques, M., Cohen, M. M., Waterham, H. R., Wanders, R. J. A., Schuldiner, M., & Zalckvar, E. (2018). Systematic mapping of contact sites reveals tethers and a function for the peroxisome-mitochondria contact. Nature Communications 2018 9:1, 9(1), 1–13. 10.1038/s41467-018-03957-8

Soukupova, M., Sprenger, C., Gorgas, K., Kunau, W. H., & Dodt, G. (1999). Identification and characterization of the human peroxin PEX3. European Journal of Cell Biology, 78(6), 357–374. 10.1016/S0171-9335(99)80078-8

Stirling, D. R., Swain-Bowden, M. J., Lucas, A. M., Carpenter, A. E., Cimini, B. A., & Goodman, A. (2021). CellProfiler 4: improvements in speed, utility and usability. BMC Bioinformatics, 22(1), 1–11. 10.1186/S12859-021-04344-9/FIGURES/6

Vallese, F., Catoni, C., Cieri, D., Barazzuol, L., Ramirez, O., Calore, V., Bonora, M., Giamogante, F., Pinton, P., Brini, M., & Calí, T. (2020). An expanded palette of improved SPLICS reporters detects multiple organelle contacts in vitro and in vivo. Nature Communications 2020 11:1, 11(1), 1–15. 10.1038/s41467-020-19892-6

Valm, A. M., Cohen, S., Legant, W. R., Melunis, J., Hershberg, U., Wait, E., Cohen, A. R., Davidson, M. W., Betzig, E., & Lippincott-Schwartz, J. (2017). Applying systems-level spectral imaging and analysis to reveal the organelle interactome. Nature, 546(7656), 162–167. 10.1038/nature22369

Varnai, P., Thyagarajan, B., Rohacs, T., & Balla, T. (2006). Rapidly inducible changes in phosphatidylinositol 4,5-bisphosphate levels influence multiple regulatory functions of the lipid in intact living cells. Journal of Cell Biology, 175(3), 377–382. 10.1083/JCB.200607116

Wang, H., Becuwe, M., Housden, B. E., Chitraju, C., Porras, A. J., Graham, M. M., Liu, X. N., Thiam, A. R., Savage, D. B., Agarwal, A. K., Garg, A., Olarte, M. J., Lin, Q., Frohlich, F., Hannibal-Bach, H. K., Upadhyayula, S., Perrimon, N., Kirchhausen, T., Ejsing, C. S., … Farese, R. V. (2016). Seipin is required for converting nascent to mature lipid droplets. ELife, 5(AUGUST). 10.7554/ELIFE.16582

Wang, H., Sreenivasan, U., Hu, H., Saladino, A., Polster, B. M., Lund, L. M., Gong, D., Stanley, W. C., & Sztalryd, C. (2011). Perilipin 5, a lipid droplet-associated protein, provides physical and metabolic linkage to mitochondria. Journal of Lipid Research, 52(12), 2159. 10.1194/JLR.M017939

Wilfling, F., Wang, H., Haas, J. T., Krahmer, N., Gould, T. J., Uchida, A., Cheng, J.-X., Graham, M., Christiano, R., Fröhlich, F., Liu, X., Buhman, K. K., Coleman, R. A., Bewersdorf, J., Farese, R. V, Walther, T. C., & Walther, T. C. (2013). Triacylglycerol synthesis enzymes mediate lipid droplet growth by relocalizing from the ER to lipid droplets. Developmental Cell, 24(4), 384–399. 10.1016/j.devcel.2013.01.013

Yang, Z., Zhao, X., Xu, J., Shang, W., & Tong, C. (2018). A novel fluorescent reporter detects plastic remodeling of mitochondria-ER contact sites. Journal of Cell Science, 131(1). 10.1242/JCS.208686/VIDEO-1

